# Exploitation of neighbouring subsoil for nutrient acquisition under annual-perennial strip intercropping systems

**DOI:** 10.1101/2022.01.09.475506

**Authors:** Eusun Han, Weronika Czaban, Dorte Bodin Dresbøll, Kristian Thorup-Kristensen

## Abstract

Little is known of how the deep root systems of perennial crops contribute to deeper and better resource use when intercropped with annuals in arable fields. Therefore, we aimed at measuring the capacity of perennial deep roots, alfalfa (*Medicago sativa* L.) and curly dock (*Rumex crispus* L.) to access the nutrient source located under the neighboring annuals at 1.0 and 2.5 m of soil depth. Alfalfa and curly dock were able to access the tracer-labelled source placed at a distance under the annual crop strips. As a result, the reliance on deeper soil layer for nutrient uptake under intercroppings became greater compared with sole-croppings. Combination of an annual cereal (winter rye) and a perennial legume (alfalfa) with contrasting root systems exhibited higher resource complementarity compared with intercroppings having similar root systems or absence of legumes. Our results demonstrated that the deep-rooted perennials when intercropped with annuals can induce vertical niche complementarity, especially at deeper soil layers. This was assumed to be due to the vertically stratified root activity between the crop components, however, the magnitude of the effects depended on choice of crop combinations, and on types of tracers. Future studies should include estimates such as relative yield total and land equivalent ratio to quantitatively determine the effects of resource acquisition under annual-perennial intercropping in arable fields.

## Introduction

Sustainable intensification leading to efficient resource use and reduced negative environmental impact is called for in modern agriculture (Thorup-Kristensen et al. 2020). Strip intercropping, the simultaneous cultivation of two or more species in a distinct row arrangement, has advantages over mono-cropping for increased crop yield. One explanation for this is the creation of more edge rows for better resource use complementarity (e.g. Maize-soybean strip intercropping; Du et al. 2018). In arable crop production, strip intercropping is often implemented using combinations of two or more annual crops such as wheat-pea, maize-cowpea and barley-pea (Hauggaard-Nielsen et al. 2001; Li et al. 2006; Streit et al. 2019). A common rhetoric has been that the crop components with different rooting depth and density do not compete for plant resources at the same soil profile, i.e., where one crop component dominates at shallower depth and the other at deeper depth (Berendse 1982; Hauggaard-Nielsen et al. 2001). Ample evidence supports belowground complementarity between crop components, leading to increased yields, i.e., Land equivalent ratio (LER)>1 (Malhi 2012). However, the depth of those studies has been rather restricted to the topsoil or shallow subsoil, whereas deep placed soil resources have been found to be important for crop nutrient supply (Han et al. 2021b).

Recent notion of growing emerging crops with deeper roots, especially perennials, has been drawing attention as an option for better exploitation from deep soil layers (Thorup-Kristensen et al. 2020). Previous studies suggest that those crops establish root systems below 2 m or even down to 4 m of soil depth (Thorup-Kristensen and Rasmussen 2015; Han et al. 2020, 2021a). Also, the recent reports on the enhanced belowground ecosystem services (e.g. increased soil organic matter accrual and nutrient acquisition) by employing perennial-based intercropping (Drinkwater et al. 2021) indicate the necessity to include perennials into the systems. Nevertheless, little is known of how strip intercropping including deep-rooted perennials affects spatial resource use in arable fields.

Increased intermingling and root density of intercropped strips has been reported, which was considered to be a main driver for greater total soil N uptake (e.g. maize-alfalfa intercropping; Zhang et al. 2013). In more extreme examples, such as in agroforestry, deep roots of tree crops showed the potential to exploit a larger soil volume by vertical penetration as well as horizontal expansion in deeper soil layers (Saize Del Rio et al. 1961; Wahid 2000; Divakara et al. 2002). However, none of those studies revealed the quantitative contribution of perennial species in terms of subsoil exploitation for nutrient uptake in strip intercropping.

In shallow soil layers, introducing new cash crops among living mulches can limit the resource acquisition capacity of the former due to the well-established root systems of the latter. Båth et al. (2008), clearly showed a reduced competition between the cash crop (white cabbage) and the perennial living mulch (birdsfoot trefoil) using root pruning at 0.2 m of soil depth. The capacity of the cash crop for N uptake increased 7-fold when the roots of perennials were pruned. Similarly, an increase in grain and stover yield of sorghum was observed when the roots of adjacent *Leucaena* hedgerows were pruned (Korwar and Radder 1994). While these studies demonstrated the greater competitive capacity of the perennials at depth, no direct evidence of perennial roots accessing the crop resources at neighboring crop strips has been reported yet, especially at depth.

Crops differ in root system architecture, and it might be the main driver determining the belowground niche complementarity under intercropping systems. As commonly adopted, cereal-legume intercropping has shown dominance of cereal over legume in terms of root depth (Hauggaard-Nielsen et al. 2001; Sun et al. 2019). Furthermore, when studied in polyculture it was found that maize, squash, and beans had distinctive rooting depths, which made the polyculture a more efficient system, especially under low nitrogen (N) availability (Postma and Lynch 2012). In some cases, such belowground interactions were attributed to the supply of the additional N to the companion crops when legumes are included (Andersen et al. 2014; Bargaz et al. 2015; Ramirez-Garcia et al. 2015). However, comparisons between intercropping systems with crop components having similar or contrasting root systems (e.g. monocot-dicot vs dicot-dicot) has rarely been carried out. Such investigation can be meaningful for areas where legume-based intercropping is not suitable, for example, in areas with high leaching risk.

Therefore, we compared the root growth and nutrient acquisition potential of three intercropping systems, winter rye-alfalfa, fodder radish-alfalfa and winter wheat-curly dock at 1 m and 2.5 m of soil depth. We hypothesize that (i) the deep roots of perennials can access the nutrients located under neighboring strips in the subsoil. As a result, (ii) the reliance on deeper soil layers for nutrient uptake increases under intercropping compared with sole-cropping. We also foresee that (iii) the resource complementarity of annual-perennial intercroppings depends on the root system architecture of each crop components as well as presence of N-fixing legume species.

## Materials and methods

### Study site

A field trial was established at the experimental station of the University of Copenhagen in Taastrup, Denmark (55 ° 40’ N; 12 ° 18’ E). The soil was classified as agrudalf (IUSS Working Group WRB 2006). The soil characteristics and weather data at the study site are available in Table 1 and Figure 1.

**Figure 1.**
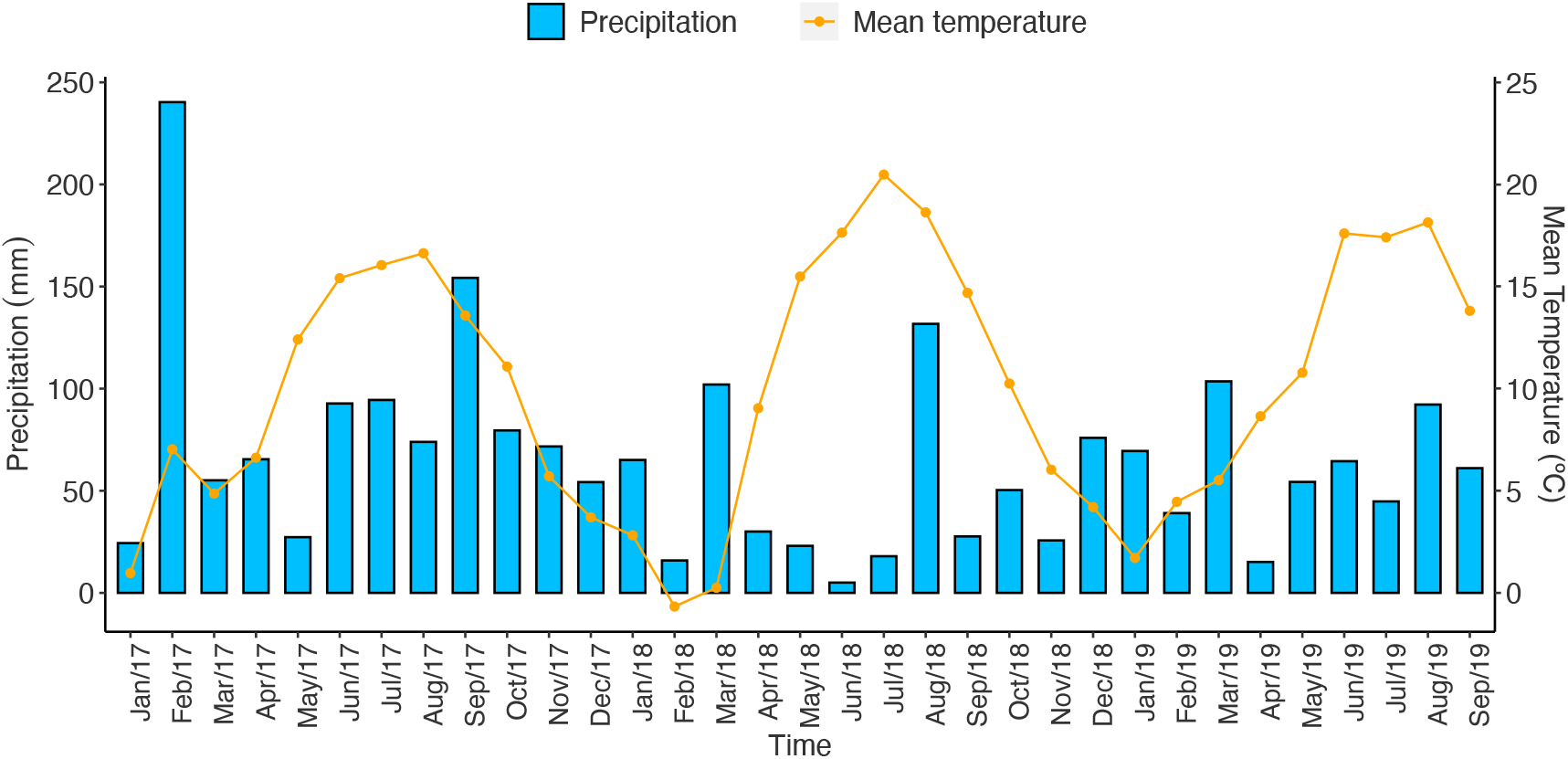
Monthly precipitation (mm) and mean temperature (°C) at the study site.

**Table 1.**
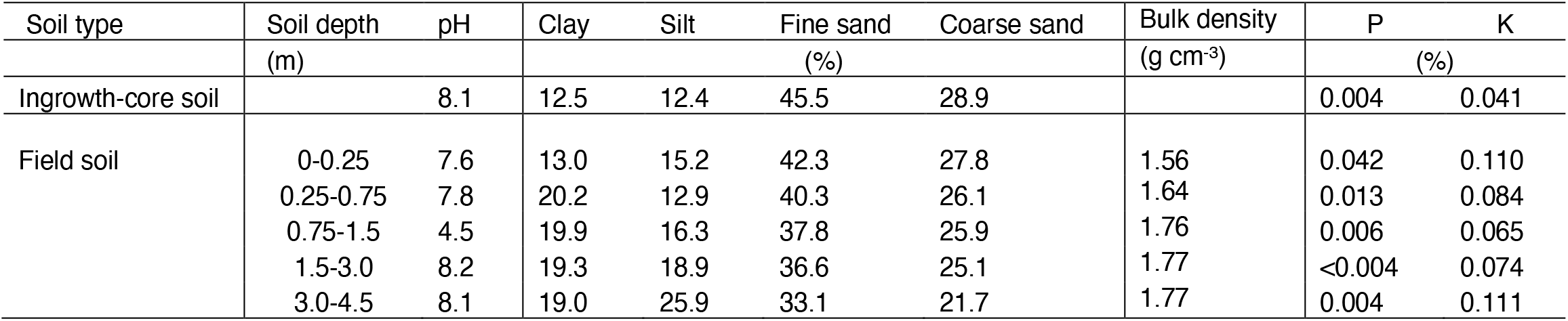
Physical and chemical soil characteristics at the study site and for the ingrowth-cores

### Core-labelling technique (CLT)

Deep root activity at the study site was measured by the core-labelling technique (CLT; Figure 2) where permanently installed metal access-tubes and insertable ingrowth-cores were used (Han et al. 2020). The access-tubes were installed at an angle of 30° vertically and have openings at 1.0 m (0.74-1.21 m) and 2.5 m (2.25-2.73 m) of soil depth. The ingrowth-cores have openings as well and can be inserted into the access-tubes to align the openings. Ingrowth-cores were packed with tracer-labelled soil and allowed a soil volume of 3931 cm^3^. ^15^NH_4_Cl (275.2 mg per ingrowth-core) and Cs2CO3 (728.3 mg per ingrowth-core) were used as nutrient tracers. ^15^N is a well-known tracer for nitrogen (N), and cesium (Cs) has been frequently used as a nutrient analogue to potassium (K). After retraction, root growth into the ingrowth cores can be determined by washing roots free of the soil. The plants growing directly above the ingrowth-core openings were used to determine tracer uptake.

**Figure 2.**
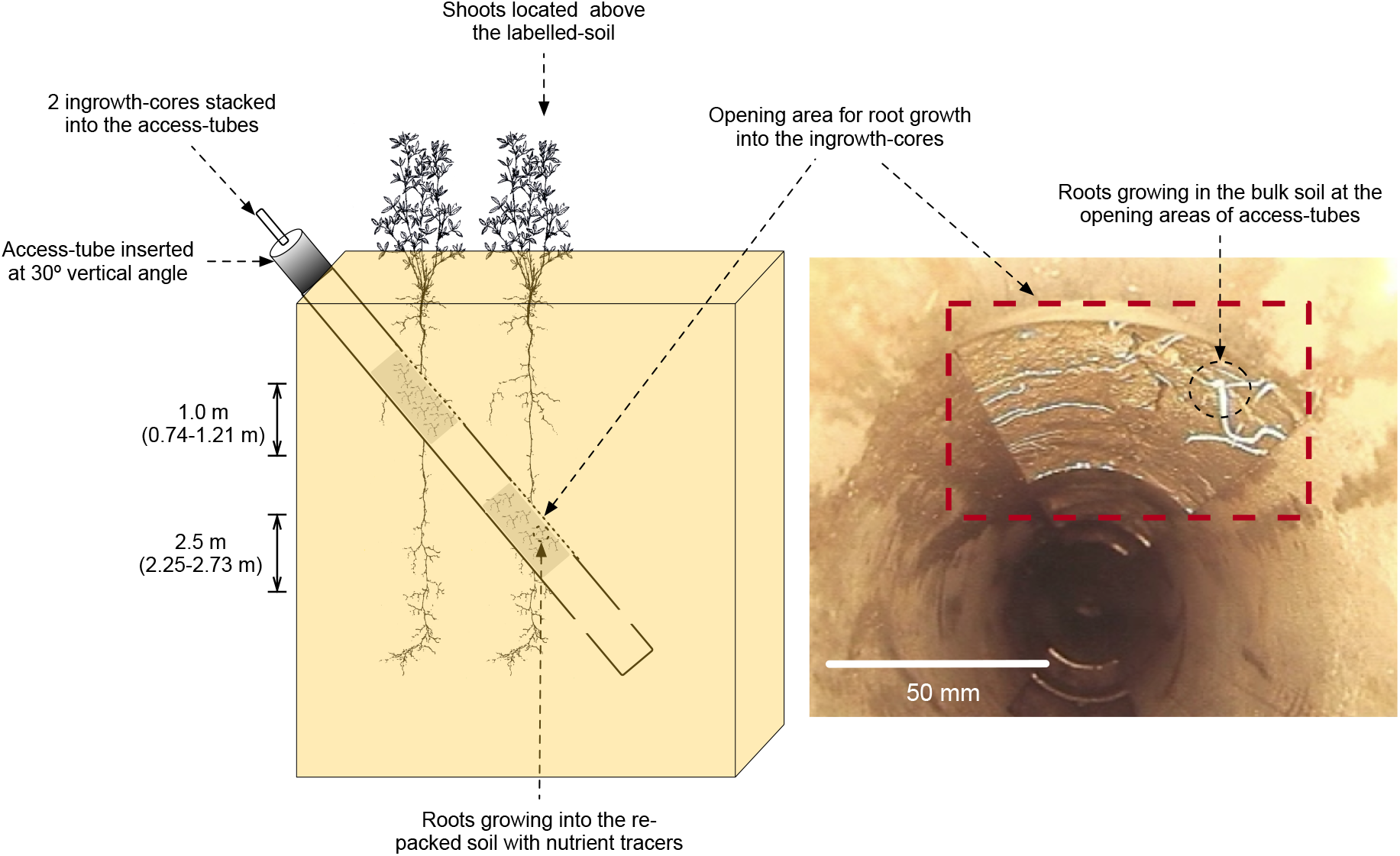
Schematic diagram of core-labelling technique (modified from Han et al. 2020)

### Experimental design

Three experiments, Exp 1 (May-July 2017), Exp 2 (Sep-Nov 2017) and Exp 3 (Apr-June 2019) were conducted. In total 3 annuals and 2 perennials were used for the study, which were winter rye (*Secale cereale* L.), fodder radish (*Raphanus sativus* L.), winter wheat (*Triticum aestivum* L.), alfalfa (*Medicago sativa* L.) and curly dock (*Rumex crispus*). In Exp 1, 2 and 3, winter rye-alfalfa, fodder radish-alfalfa and winter wheat-curly dock combinations were tested respectively. Sowing dates, seeding density and fertilizer application rates are shown in Table 2.

**Table 2.** Experimental design, sowing dates, seeding density and fertilizer application rate

| Year | Month | Experiment | Crop components | Sowing dates | Seeding density (kg ha<sup>-1</sup>) | Fertilizer application (kg N ha<sup>-1</sup>) |
| --- | --- | --- | --- | --- | --- | --- |
| 2017 | May-July | Exp 1 | Alfalfa | 09/09/2015 | 25 | - |
|  |  |  | Winter rye | 24/08/2016 | 115 | 60 |
| 2017 | Sep-Nov | Exp 2 | Alfalfa | 09/09/2015 | 25 | - |
|  |  |  | Fodder radish | 31/07/2017 | 50 plant m<sup>-2</sup> | 60 |
| 2019 | Apr-June | Exp 3 | Curly dock | 03/05/2017 | 4 | 90 |
|  |  |  | Winter wheat | 28/09/2018 | 234 | 100 |

The annual strips were 1.5×10 m while the perennial strips were 3×10 m long. Plants in the middle of the strip was treated as sole-crops, while plants at the edge of the strip was treated as intercrops. This made it possible to create four treatments; sole-cropped and intercropped annuals, and sole-cropped and intercropped perennials (Figure 3).

**Figure 3.**
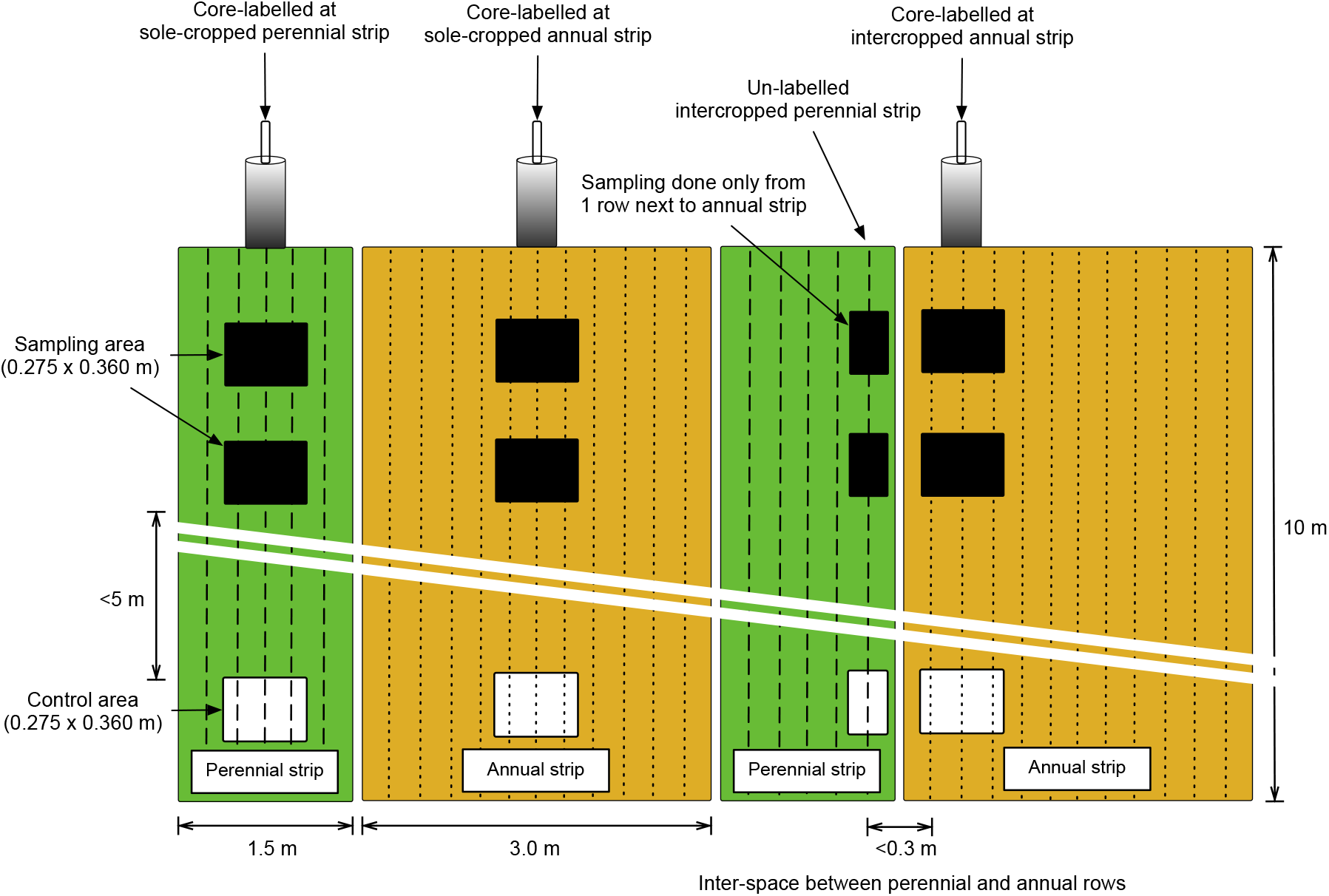
Schematic diagram of strip intercropping design seen from above

The ingrowth-cores were filled with tracer labelled soil and inserted into the access-tubes at the beginning of the experimental period and retracted at cessation of the experiment. The distance between annual strips and perennial strips was set to approximately 0.3 m. However, due to the sowing and tillage management, some of the strips were up to 0.5 m apart. The larger distance did not affect the results. Each treatment contained six replicates.

Our main objective was to measure the capacity of perennials deep roots to reach the neighboring soil profile occupied by the annuals. The intercropped annuals were target-labelled by inserting soil samples with nutrient tracers directly under the row, whereas intercropped perennials were not labelled directly, but grown at distance (<0.3 m) from the nutrient tracers.

### Root sampling and measurement

Root samples were acquired from the ingrowth-cores inserted at sole-cropped and intercropped annuals and sole-cropped perennials. As explained above, ingrowth-cores were not inserted directly below the intercropped perennials, why no root samples were collected below these treatments. The root samples from intercropped annuals were assumed to contain roots from both crops.

Roots were washed free of soil and the obtained root samples were photo-scanned using an Epson Perfection V700 resulting in root images in **TIFF** format (600 dots per inch; DPI). Using the ‘WinRHIZO Pro’ (Version 2016c, 32 Bit) software, the root images were analyzed for root length (cm), which was then calculated further to rootlength density (cm cm^-3^) based on the soil volume. Upon analysis, minimum surface area was set to 2 cm^2^, and length to width ratio of the root objects considered was 2. Medium image smoothening was chosen for noise removal. For each root image, roots were divided into four root-size classes, ≤0.1 mm, 0.11-0.20 mm, 0.21-0.5 mm and ≥ 0.5 mm based on Reinhardt and Miller (1990), from which, the proportion of root length (%) per root-size class was calculated.

### Shoot sampling and measurement

Shoot samples were collected from an area of 0.275 m x 0.36 m vertically above where the ingrowth-cores were inserted at 1.0 and 2.5 m of soil depth (Figure 3). Shoot samples from the row of the perennial strips adjacent to the annual strips were additionally collected as an un-labelled intercropped perennial strip.

For all three experiments, shoot samples were collected 4 and 8 weeks after ingrowth-cores were inserted for measurements of ^15^N and Cs. At sampling week 4, 10 shoots per sampling area were randomly selected. For alfalfa we collected the young shoots from the three top branches of the plants. The curly dock and fodder radish were still at early vegetative stages; therefore, we chose any uncurled 10 leaves from the plants. The young shoots from the cereals were collected up to the 1st internode from the top including the spikes when present. At week 8, the entire biomass from the sampling area was collected. For intercropped perennials, biomass was collected only from the single row directly adjacent to the annual strips.

The collected samples were oven-dried at 85°C for 48 hours, and finely ground for further analysis. Stable isotopic ratios of N (δ^15^N) were measured at the Stable Isotope Facility, UC Davis, using a ThermoScientific GasBench-Precon gas concentration system interfaced to a TheromScintific Delta V Plus isotope-ratio mass spectrometer (Bremen, Germany). Upon analysis of Cs, the samples were microwave-digested in nitric acid (70 %). Sample digests were analyzed by Inductively Coupled Plasma Sector Field Mass Spectrometry (ICP-SFMS, ELEMENT XR, ThermoScientific, Bremen, Germany) using a combination of internal standardization and external calibration.

### Calculations

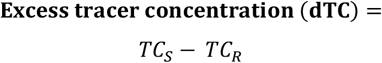

Excess tracer concentration (dTC) was calculated based on Hoekstra et al. (2014), where TCs denotes tracer concentration at sampling area, and TC_R_ at control area.

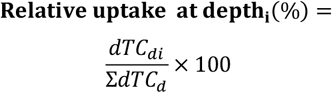

Relative uptake (%) was calculated based on Da silva et al. (2011). The dTC values obtained at depth i (dTC_di_) were divided by the sum of dTC at all soil depth (∑dTC_d_) and multiplied by 100 to determine the percentage (%).

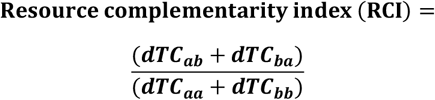

Relative yield total (RYT) was suggested by De wit and den Bergh (1965) as a measure of resource use between the intercropped and sole-cropped crop components. We modified the equation to reveal the resource complementarity of strip intercropping over sole cropping, where sum of dTC_ab_ and dTC_ba_ was divided by the sum of dTCaa and dTCbb, called resource complementarity index (RCI). A RCI value higher than unity indicates that the strip intercropping resulted in higher resource use compared with sole cropping.

### Statistical analysis

R version 3.4.1(R Development Core Team 2019) was used for statistical analysis. The package lme4 (Bates et al. 2013) was used for linear mixed-effects model analysis (Pinheiro and Bates 2000). For RLD, dTC and RCI, crop strip and soil depth were treated as one fixed factor. For RLD, ingrowth-core ID was used as a random factor. For dTC and RCI, sampling time and plot were used set as random factors. Proportion of root length was tested against crop strip at each depth. Log-transformed variables were used for statistical analysis. Main effects and interactions were tested for significance (P≤0.05) based on the approximated degrees of freedom calculated by the package ImerTest (Kuznetsova et al. 2015). Estimates and standard errors were back transformed using the delta method and reported on the original scale for dTC and RCI, whereas mean and standard errors were shown for RLD, proportion of root length and relative uptake. Post-hoc tests were performed by multiple comparisons (Tukey HSD; P≤0.05) using the package Multcomp (Hothorn et al. 2019).

## Results

### Excess tracer concentration (dTC)

Calculation of the excess tracer concentrations (dTC) revealed net tracer uptake at sole cropped as well as at intercropped strips. Most importantly, the un-labelled intercropped perennials, next to the intercropped annuals, also resulted in positive dTC values in all three experiments. However, in some cases the used tracers (^15^N and Cs) led to inconsistent results.

In Exp 1, the sole-cropped and intercropped winter rye at 1.0 m revealed greater ^15^N uptake compared with other treatments (Figure 4a). There was a clear trend of depthwise difference in ^15^N uptake within the treatments except the un-labelled alfalfa strips. In Exp 2, we noticed that the overall ^15^N uptake by each crop components was substantially smaller compared with those observed in the other two experiments (Figure 4b). Similar results were found in Exp 3, where ^15^N uptake was dominantly shown at 1.0 m (Figure 4c).

**Figure 4.**
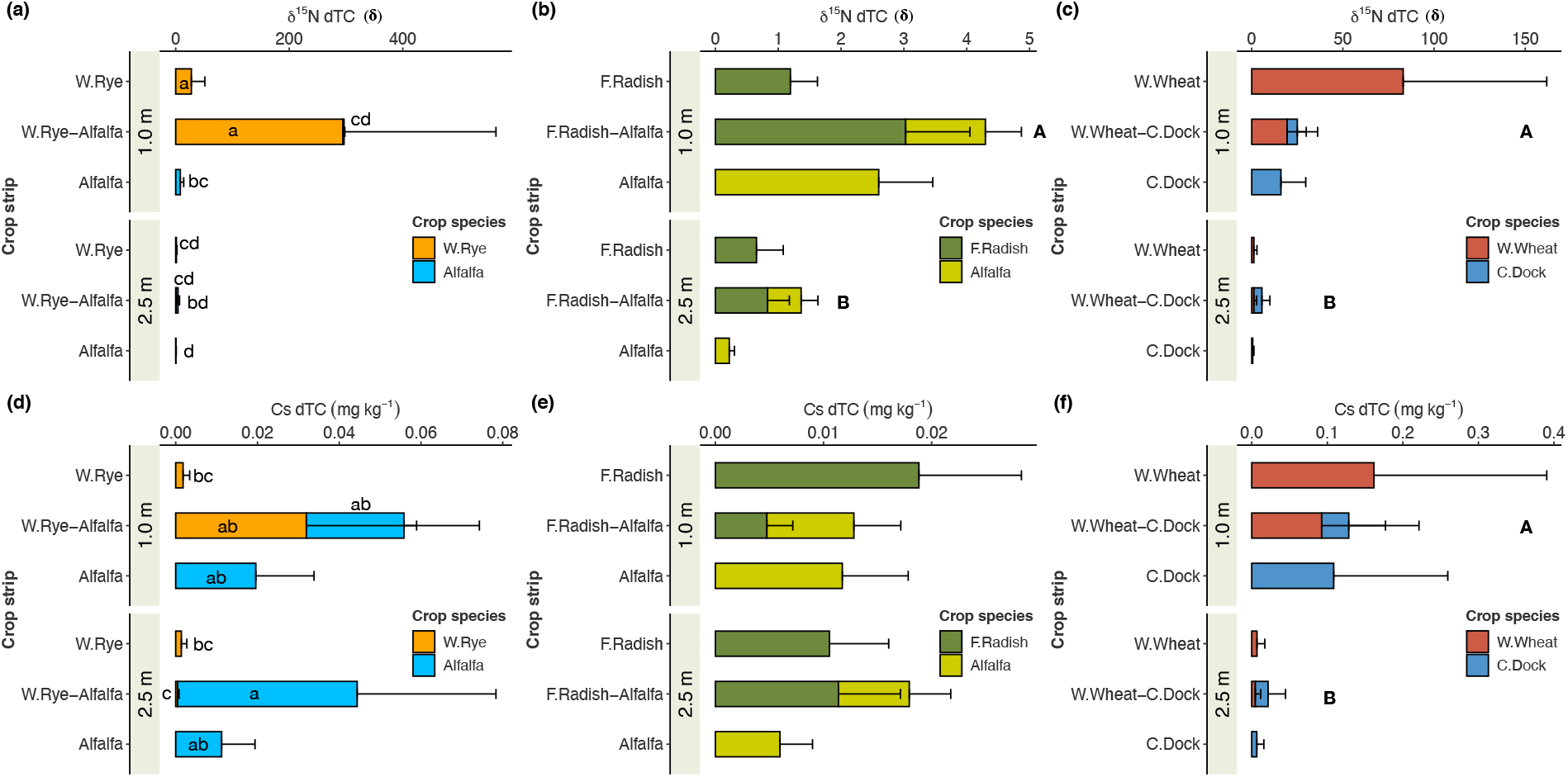
Excessive tracer concentration (dTC) affected by crop species and soil depth (1.0 and 2.5 m) exhibited in Exp 1 (a,d), 2 (b,e) and 3 (c,f) Small and capital letters indicate significant differences between the crop species and soil depth, respectively (Tukey HSD; P≤0.05). Mixed-effects model was run against two fixed factors (crop species and soil depth). Log-transformed variables used were back-transformed and the Estimates and SE are shown here (n=6).

Similar measurements were carried out with the tracer Cs. In Exp 1, substantial Cs uptake was noted for the un-labelled alfalfa at 2.5 m – which was significantly different to Cs uptake by sole-cropped winter rye at both depths, and intercropped winter rye at 2.5 m. Other treatments exhibited intermediate differences (Figure 4d). In Exp 2 (Figure 4e) and 3 (Figure 4f), there were no significant differences between the crop species. A depth-wise difference in Cs uptake was noted in Exp 3.

### Relative uptake

Based on the excessive tracer concentrations, we calculated the relative uptake, which is the proportion of tracer uptake from respectively 1.0 and 2.5 m soil depth of the total uptake. The results indicate that the tracer uptake occurred more actively at 1.0 m for sole-cropped strips compared with the tracer uptake at 2.5 m. Except for one occasion in Exp 3, the tracer uptake from the intercropped perennials were similar between the soil depths.

Calculated from δ^15^N dTC the sole-cropped winter rye, intercropped winter rye and sole-cropped alfalfa in Exp 1 exhibited greater relative uptake at 1.0 m compared with 2.5 m. The difference became insignificant for intercropped alfalfa (Figure 5a). Similar results were found in Exp 2, in which, all other treatments except the intercropped alfalfa showed significantly greater relative uptake from 1.0 m. In Exp 3, all crop strips exhibited greater relative uptake from 1.0 m including the intercropped curly dock.

**Figure 5.**
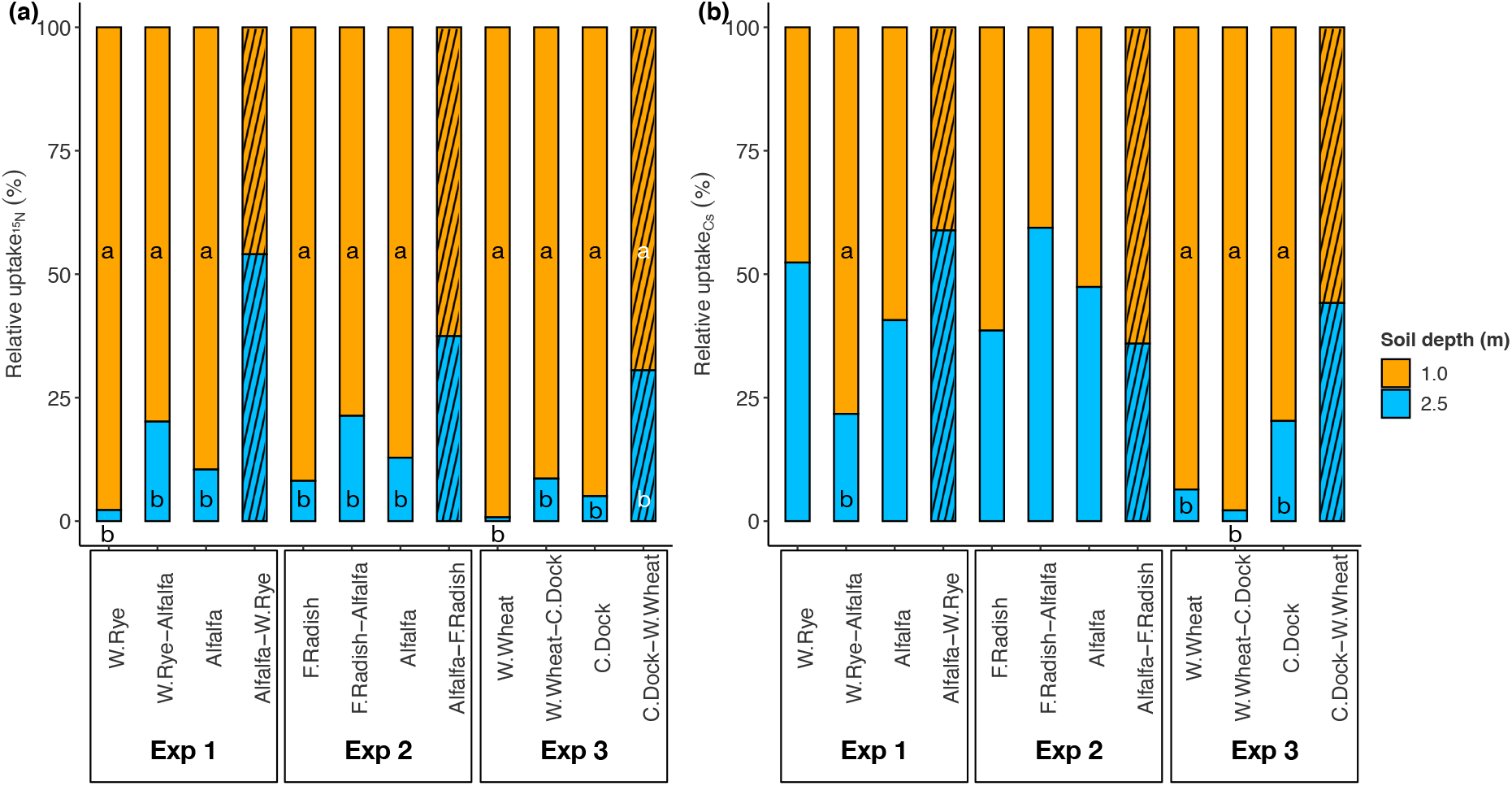
Relative uptake (%) between soil depth (1.0 m and 2.5 m) of each crop strips calculated from δ^15^N dTC (a) and Cs dTC (b). Mixed-effects model was run against soil depth. Small letters indicate significant differences between the soil depth within the crop strips (Tukey HSD; P≤0.05). Mean and SE are shown here (n=6).

The depth-wise difference in relative uptake of Cs dTC was less apparent than δ^15^N dTC (Figure 5b). In Exp 1, only the intercropped winter rye strips exhibited greater relative uptake at 1.0 m. No depth-effects were present in Exp 2, meaning, the proportion of Cs dTC between the two depth-levels was statistically the same for all the crop strips. In Exp 3, greater relative uptake at 1.0 m was shown for the sole-cropped wheat, curly dock, and intercropped wheat, whereas no such difference was found at the intercropped curly dock strips.

### Resource complementarity index

We have determined the resource complementarity index (RCI) as a substitute measure for the generally used relative yield potential (RYT). When measured ^15^N uptake, winter rye-alfalfa and fodder radish-alfalfa intercropping revealed RCI over unity at both soil depths – meaning the intercropping was more efficient in resource use compared with the sole-cropping. When measured by Cs uptake, winter rye-alfalfa intercropping was found to be a more efficient system compared with their corresponding sole-cropping at both soil layers. A similar finding was seen in winter wheat-curly dock intercropping at 2.5 m.

Comparisons of RCI between the soil depths and intercropping treatments were carried out. Regardless of soil depth, RCI of winter rye-alfalfa intercropping measured based on ^15^N uptake was greater than RCI of winter wheat-curly dock intercropping (Figure 6a). Fodder radish-alfalfa intercropping exhibited an intermediate RCI. Across the intercropping treatments, RCI measured from ^15^N uptake was greater at 2.5 m compared with RCI at 1.0 m. RCI measured from Cs uptake did not reveal a difference between the soil depths (Figure 6b). Between the different intercropping combinations, winter rye-alfalfa intercropping revealed the greater RCI over other intercropping treatments across the depths.

**Figure 6.**
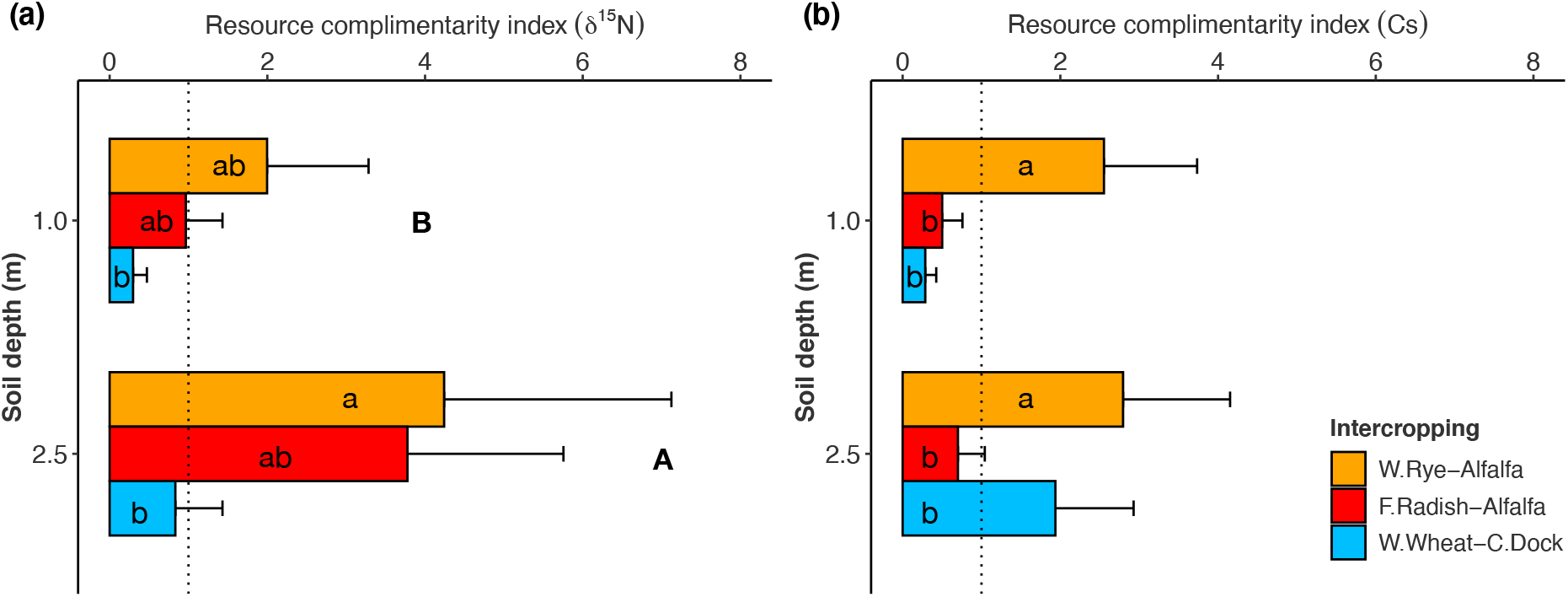
Resource complementarity index of ^15^N (a) and Cs (b) affected by intercropping (Winter rye-Alfalfa, Fodder radish-Alfalfa and Winter wheat-Curly dock) and soil depth (1.0 and 2.5 m). Small and capital letters indicate significant differences between intercropping and soil depth, respectively (Tukey HSD; P≤0.05). Log-transformed variables used were back-transformed and estimates and SE are shown here (n=6).

### Root growth

We measured the root-length density (RLD; cm cm^-3^) of the newly grown roots into the ingrowth-cores containing the labelled soil (Figure 7) and did not find significant evidence that crop species and intercropping affected the new root growth within 8 weeks of time. However, a depth-wise difference in RLD was apparent in Exp 1 and 3. Nevertheless, there was a trend showing that sole-cropped perennials grew fewer fine roots compared with other treatments at 1.0 m during the incubation time, although not significantly.

**Figure 7.**
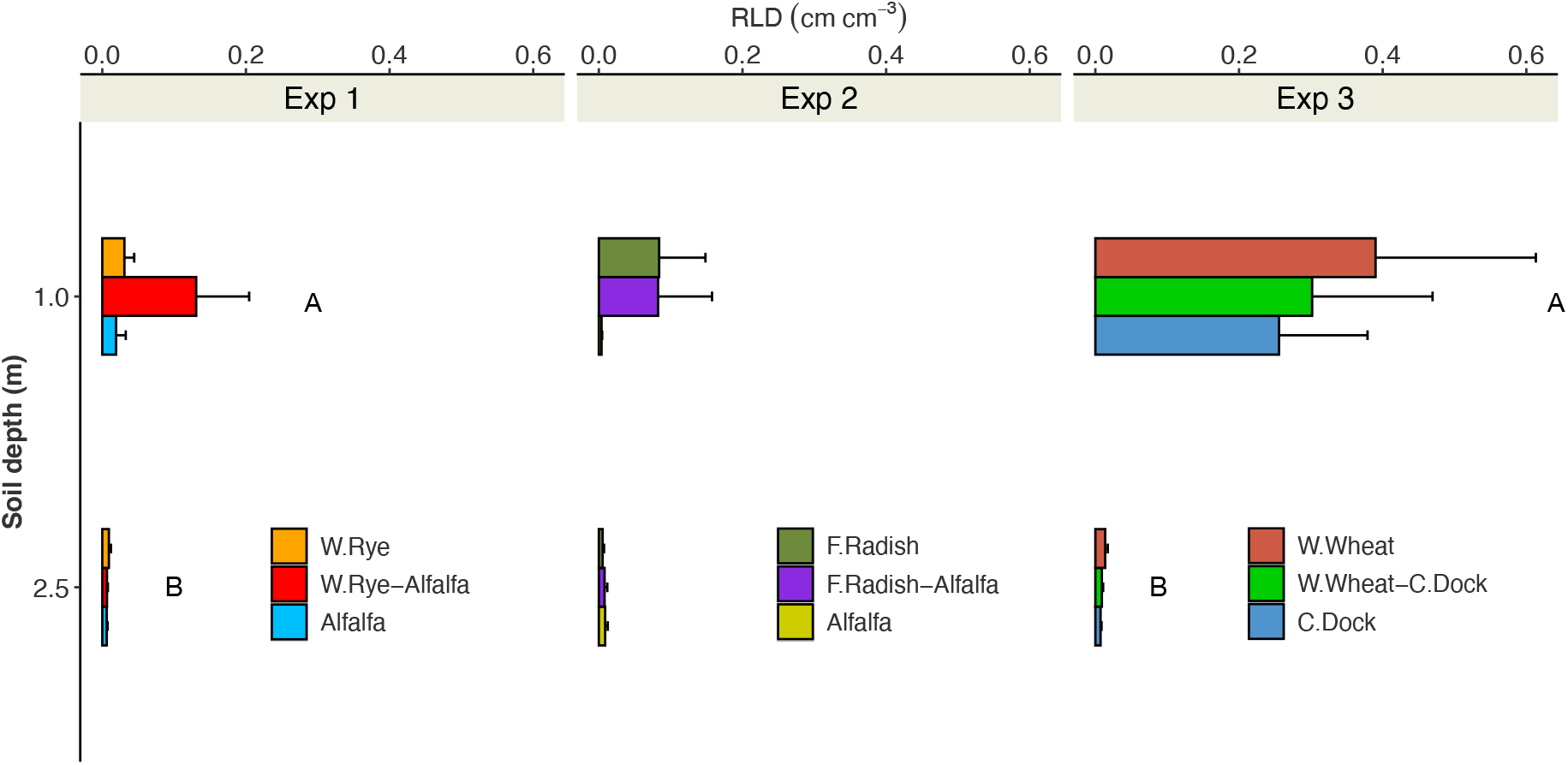
Root-length d ensity(cmcm^-3^) affected by crop strips (sole-cropped annual, intercropped annual and sole-cropped perennial strips) and soil depth (1.0 and 2.5 m) exhibited in Exp 1, 2 and 3 (a-c). Mixed-effects model was run against one fixed factor combining crop strip and soil depth with a log-transformed variable. Small letters indicate significant differences between all the treatments (Tukey HSD; P≤0.05). Mean and SE are shown here (n=6).

Using the root samples acquired from the ingrowth-cores, we calculated the proportion of root length (%) in four root-size classes (≤0.1,0.11-0.20, 0.21-0.50 and ≥0.51 mm). We expected that the analysis might describe the root mixture of annuals and perennials when intercropped as its morphology differs from the monocultured crop components. In Exp 1, the proportion of roots with a diameter of 0.11 −0.20 mm at the sole-cropped winter rye strips was larger than intercropped winter rye, meanwhile sole-cropped alfalfa showed a moderate difference (Figure 8). In Exp 3, the sole-cropped winter wheat exhibited a greater proportion of larger roots (≥0.51 mm) compared to sole-cropped curly dock. No changes in root size-classes were found in Exp 2.

**Figure 8.**
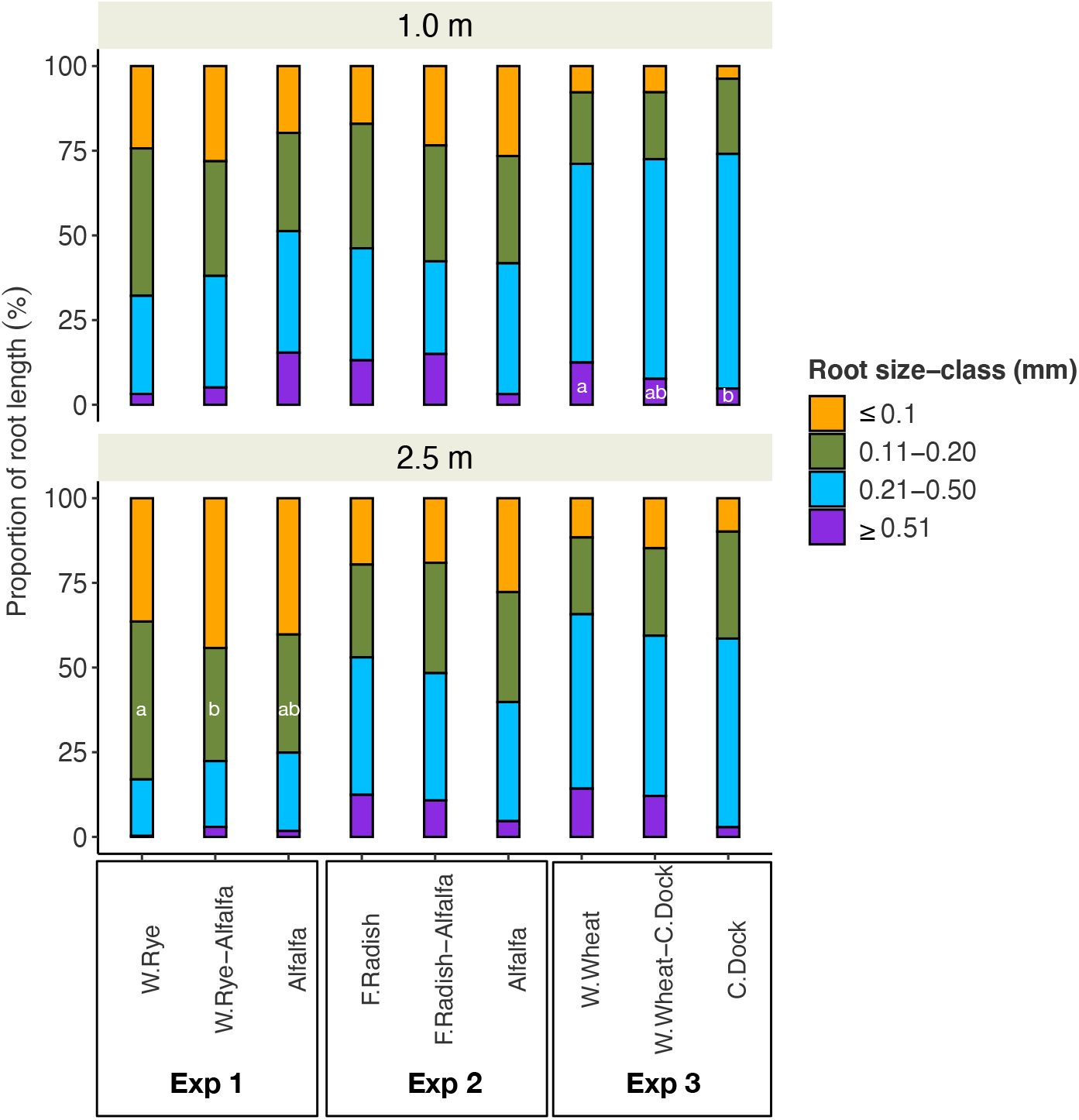
Proportion of root length (%) affected by crop strip (sole-cropped annual, intercropped annual, sole-cropped perennial) in Exp 1, 2 and 3 at 1.0 m and 2.5 m. Mixed-effects model was run against one fixed factor (crop strip) with a log-transformed variable. Small letters indicate significant differences between the crop strips within the root-size class (Tukey HSD; P≤0.05). Mean and SE are shown here (n=6).

## Discussion

### Subsoil accessibility in strip intercropping

We acquired clear evidence that the deep-rooted perennials in the strip intercropping were able to access the tracer sources located at the neighboring annual strips, and thereby confirmed the hypothesis on belowground complementarity mediated by perennial deep roots. Except for ^15^N uptake in Exp 1, the tracer uptake by the unlabelled perennials was at similar levels to the labelled crop strips. In an extreme case, the tracer uptake by alfalfa, the un-labelled perennial, was even more substantial than the labelled treatments (Cs uptake at 2.0 m in Exp1).

Similar observations were shown in annual row crops at shallower soil depth. For example, wheat extended its root systems under the root zone of maize when intercropped and measured to 60 cm of soil depth (Li et al. 2006). The row distance between the two species was 30 cm, which was similar to the row distance in our study. Zhang et al. (2013) concluded that maize root density increased both vertically and laterally when intercropped with alfalfa at 30 cm distance as well. The new aspect of our results is that such horizontal root growth by deeper rooted crop species in intercropping still can prevail up to 2.5 m of soil, which can be meaningful in the regions where topsoil nutrient availability is low.

Based on the observation of root depth and distribution in the bulk soil (data not shown), perennial crops tended to exhibit deeper rooting depth and often greater root density than the annuals. In our study, however, the measured RLD from ingrowth-cores did not reveal significant differences between the crop strips. Effects of intercropping can rather be found from the root morphological analysis from our study, which exhibited significant changes. The higher proportion of 0.11-0.20 mm-sized roots were found in sole cropped winter rye compared to intercropped winter rye in Exp 1, which indicate that the ingrowth-cores below the intercropped winter rye contained a mixture of roots from both crop species. Similarly, in Exp 3, the intercropped winter wheat showed a shift in large root class sizes (≥0.51 mm) from sole cropped winter wheat and curly dock. Also, the intercropped strips tended to exhibit an intermediate root growth between the sole-cropped strips of annuals or perennials, except for winter rye-alfalfa intercropping. This is in line with a previous study by Ramirez-Garcia et al. (2015) who found that root intensity (crosses m^-1^) of barley-vetch intercropping was in-between barley and vetch monocrops at the 1.2 −1.6 m soil layer observed on rhizotrons. Therefore, we conclude that the growth of fine roots into the ingrowth-cores partially represents the effects of intercropping.

### Deeper accessibility

The belowground complementarity was found to be more substantial at deeper soil layers, thereby confirming the second hypothesis. The labelled crop strips clearly showed greater relative uptake of tracers at 1.0 m, whereas the un-labelled perennials did not exhibit such differences with one exception in Exp 3. These results strongly indicate the potential of arable subsoil under 1 m to contribute to plant nutrient uptake under intercropping systems.

This is also in line with previous reports, where the root fractions of intercropped wheat and bean increased at deeper soil layers compared to sole-cropped plants (Streit et al. 2019). Another example of perennials is the bamboo-teak root interactions shown by Divakara et al. (2002). When teak trees were labelled with ^32^P-isotope, the root activity was found to be greater at subsoil layers (0.5 m) compared with the topsoil layers. The authors speculated that the dicot (teak) root system was forced to grow deeper due to the co-proliferating monocot (bamboo). Similarly, Mommer et al. (2010) have found an overyielding of belowground biomass under grass mixtures when investigated to 6.6 m of soil depth. The previous studies concluded that the increased root mass density was driven by one species’ enhanced root investment. Our approach cannot confirm such species-wise interactions. However, our results on tracer concentrations from winter rye-alfalfa and fodder radish-alfalfa intercropping, showing a higher uptake than from perennials grown as sole crops, might describe the stimulated root growth of the annuals by the neighboring perennial crops.

### Resource complementarity between intercroppings

The results on the resource complementarity index (RCI) were variable depending on the intercropping systems. We assumed that the differences in root system architecture between the intercropped crops can affect belowground complementarity and thereby resource uptake. This was not fully supported by our results as only moderate differences between the intercropping in terms of RCI was found. Although winter rye-alfalfa intercropping tended to lead to a greater complementarity than fodder radish-alfalfa intercropping, the results were not conclusive to support our hypothesis (iii).

Nevertheless, there was a more apparent contrast visualized between the intercropping systems with and without legumes in this study, i.e., winter rye-alfalfa vs. winter wheat-curly dock intercropping. At 1.0 m of soil depth the legume-involved intercropping resulted in a greater resource complementarity index (Cs). Moreover, in most cases, when the intercropping involved alfalfa in this study, the resource complementarity index tended to be greater. Therefore, it is speculated that the neighboring annual species’ root growth was enhanced by the increased soil N status (Ramirez-Garcia et al. 2015). Also, the overall soil fertility and other ecosystem services provided by the legume species might have been accounted for in these effects. In a recent study it was shown that the availability of P from the plant-extractable pool was doubled when maize was intercropped with *Desmodium* spp., a perennial legume (Drinkwater et al. 2021). The authors suspected that the elevated soil organic matter increased the accrual of the added P fertilizer in that system. In our case, it might be related to the N effects of legume influencing the root growth of the neighboring crop. It has been shown that the nodule formation of alfalfa becomes deeper at least up to 80 cm of soil depth as the plants mature (1 year < 2 years < 3 years; Li et al. 2013). At the time of Exp 1 and 2, alfalfa was over 2 years old, and might have been active in N fixation in the subsoil layers which might have led to the increased root growth of winter rye. This is known as root foraging-as claimed by Hutchings and John (2003). However, we have not measured soil N content as affected by the N fixation at the depth levels we have tested for belowground interactions, which should be followed by further studies.

### Intercropping with perennials in practice

Cultivation of perennials in crop land can have several advantages such as increased soil organic matter (de Oliveira et al. 2019), increased water availability (Gaiser et al. 2012) and subsoil amelioration with deep penetration capacity (Huang et al. 2020). Also, in contrast to agroforestry systems, intercropping with perennial crops can have advantages such as reduced aboveground competition for light, which can be beneficial for the commonly grown main crops. When soybean and peanut were intercropped with apple trees, photosynthetically active radiation (PAR) and net photosynthetic rate (NPR) decreased when reducing distance between the crop components from 2.5 to 0.5 m (Gao et al. 2013).

We did not quantify the relative yield total (RYT) as suggested by De wit and den Bergh (1965). Our approach mainly aimed at determining the change in tracer concentrations of different treatments rather than quantifying the overyielding per area. The modified equation, relative complementarity index (RCI), however, has resulted in greater index values than unity of all three intercropping treatments. It indicates that placing perennial crop species near the main annual crops was beneficial in terms of spatial resource use.

Nevertheless, our study demonstrated the potential use of deep-rooted perennials in intercropping systems, and justification of the approach in practice still requires further confirmation on crop yield and other farm economic parameters. Intercropping between perennial and annual crops can be justified when the former yields commercially viable products such as fodder (e.g. alfalfa), grains (intermediate wheatgrass) or any other form of generating income for farmers (Schroth 1998). Another aspect not covered in this study is the topsoil layers, where practical tillage is usually implemented for crop production. The pre-established root systems of perennials can also give a temporal advantage over newly sown annuals in terms of resource competition (Schroth 1998). However, this can pose pressure on the annuals to develop their root systems. In this case, soil tillage on the plots where the annuals are to be sown can shift the competitive balance by removing the superficial roots of the perennials which was shown previously by Båth et al. (2008) as well as Korwar and Radder (1994) by root pruning.

## Conclusions

Our results demonstrated that the deep-rooted perennials when intercropped with annuals can induce vertical niche complementarity, especially at deeper soil layers. This was invoked by the extended horizontal root growth by the deep roots toward the neighboring subsoil. The magnitude of the effects depended on choice of crop combinations, and on types of tracers used. Future studies should estimate the practical parameters such as relative yield total and land equivalent ratio to quantitatively determine the effects of resource acquisition under annual-perennial intercropping in arable fields.

## Authors’ contribution

All authors contributed to manuscript writing. EH designed and conducted the research and prepared the manuscript. WC contributed to early conceptualization of this manuscript. KTK and DBD provided the funding and resources.

## Acknowledgements

We are grateful to **Alan Hansen** for co-designing and building the access-tubes and ingrowth-cores. Special thanks to **Paolo Mucci** who provided assistance for early testing and validating core-labelling technique (CLT) as used in this study. We would like to thank the research technicians, especially **Aymeric d’Herouville** and **Jason Allen Team** at Plant Facilities and the Workshop of the University of Copenhagen for their effortless support.

## References

Andersen SN, Dresbøll DB, Thorup-Kristensen K (2014) Root interactions between intercropped legumes and non-legumes-a competition study of red clover and red beet at different nitrogen levels. Plant and Soil 378:59–72. https://doi.org/10.1007/s11104-013-2014-4

Bargaz A, Isaac ME, Jensen ES, Carlsson G (2015) Intercropping of Faba Bean with Wheat Under Low Water Availability Promotes Faba Bean Nodulation and Root Growth in Deeper Soil Layers. Procedia Environmental Sciences 29:111–112. https://doi.org/10.1016/j.proenv.2015.07.188

Bates D, Maechler M, Bolker B (2013) lme4: Linear mixed-effects models using S4 classes, R package

Båth B, Kristensen HL, Thorup-Kristensen K (2008) Root pruning reduces root competition and increases crop growth in a living mulch cropping system. Journal of Plant Interactions 3:211–221. https://doi.org/10.1080/17429140801975161

Berendse F (1982) Competition between plant populations with different rooting depths III. Field experiments. Oecologia 53:50–55

Da Silva EV, Bouillet JP, De Moraes Gonçalves JL, et al (2011) Functional specialization of Eucalyptus fine roots: Contrasting potential uptake rates for nitrogen, potassium and calcium tracers at varying soil depths. Functional Ecology 25:996–1006. https://doi.org/10.1111/j.1365-2435.2011.01867.x

de Oliveira G, Brunsell NA, Crews TE, et al (2019) Carbon and water relations in perennial Kernza (Thinopyrum intermedium): An overview. Plant Science 110279. https://doi.org/10.1016/j.plantsci.2019.110279

de Wit CT, van den Bergh JP (1965) Competition between herbage plants. Netherlands Journal of Agricultural Science 13:212–221

Divakara B, Balachandran P V, Kumar BM (2002) Bamboo hedgerow systems in Kerala, India: Root distribution and competition with trees for phosphorus. Agroforestry Systems 103:239–248. https://doi.org/10.1023/A:1010730314507

Drinkwater LE, Midega CAO, Awuor R, et al (2021) Perennial legume intercrops provide multiple belowground ecosystem services in smallholder farming systems. Agriculture, Ecosystems and Environment 320:107566. https://doi.org/10.1016/j.agee.2021.107566

Du JB, Fu HT, Gai JY, et al (2018) Maize-soybean strip intercropping: Achieved a balance between high productivity and sustainability. Journal of Integrative Agriculture 17:747–754. https://doi.org/10.1016/S2095-3119(17)61789-1

Gaiser T, Perkons U, Küpper PM, et al (2012) Evidence of improved water uptake from subsoil by spring wheat following lucerne in a temperate humid climate. Field Crops Research 126:56–62. https://doi.org/10.1016/j.fcr.2011.09.019

Gao L, Xu H, Bi H, et al (2013) Intercropping Competition between Apple Trees and Crops in Agroforestry Systems on the Loess Plateau of China. PLoS ONE 8:1–8. https://doi.org/10.1371/journal.pone.0070739

Han E, Dresbøll DB, Thorup-Kristensen K (2020) Core-labelling technique (CLT): a novel combination of the ingrowth-core method and tracer technique for deep root study. Plant Methods 16:84. https://doi.org/10.1186/s13007-020-00622-4

Han E, Dresbøll DB, Thorup-Kristensen K (2021a) Tracing deep P uptake potential in arable subsoil using radioactive 33P isotope. Plant and Soil. https://doi.org/10.1007/s11104-021-05178-3

Han E, Li F, Perkons U, et al (2021b) Can precrops uplift subsoil nutrients to topsoil? Plant and Soil 463:329–345. https://doi.org/10.1007/s11104-021-04910-3

Hauggaard-Nielsen H, Ambus P, Jensen ES (2001) Temporal and spatial distribution of roots and competition for nitrogen in pea-barley intercrops - a field study employing P-32 technique. Plant and Soil 236:63–74

Hoekstra NJ, Finn JA, Buchmann N, et al (2014) Methodological tests of the use of trace elements as tracers to assess root activity. Plant and Soil 380:265–283. https://doi.org/10.1007/s11104-014-2048-2

Hothorn T, Bretz F, Westfall P, Heiberger RM (2019) Package “multcomp”

Huang N, Athmann M, Han E (2020) Biopore-Induced Deep Root Traits of Two Winter Crops. Agriculture 10:634. https://doi.org/10.3390/agriculture10120634

Hutchings MJ, John EA (2003) Distribution of Roots in Soil, and Root Foraging Activity. pp 33–60

Korwar GR, Radder GD (1994) Influence of root pruning and cutting interval ofLeucaena hedgerows on performance of alley croppedrabi sorghum. Agroforestry Systems 25:95–109. https://doi.org/10.1007/BF00705670

Kuznetsova A, Brockhoff PB, Christensen RHB (2015) Package “lmerTest.” R package version

Li L, Sun J, Zhang F, et al (2006) Root distribution and interactions between intercropped species. Oecologia 147:280–290. https://doi.org/10.1007/s00442-005-0256-4

Malhi SS (2012) Improving crop yield, N uptake and economic returns by intercropping barley or canola with pea. Agricultural Sciences 03:1023–1033. https://doi.org/10.4236/as.2012.38124

Mommer L, van Ruijven J, de Caluwe H, et al (2010) Unveiling below-ground species abundance in a biodiversity experiment: A test of vertical niche differentiation among grassland species. Journal of Ecology 98:1117–1127. https://doi.org/10.1111/j.1365-2745.2010.01702.x

Pinheiro J, Bates D (2000) Mixed-Effects Models in S and S-PLUS. Springer, New York

Postma JA, Lynch JP (2012) Complementarity in root architecture for nutrient uptake in ancient maize/bean and maize/bean/squash polycultures. Annals of botany 110:521–534. https://doi.org/10.1093/aob/mcs082

R Development Core Team (2019) R: A language and environment for statistical computing.

Ramirez-Garcia J, Martens HJ, Quemada M, Thorup-Kristensen K (2015) Intercropping effect on root growth and nitrogen uptake at different nitrogen levels. Journal of Plant Ecology 8:380–389

Reinhardt DR, Miller RM (1990) Size classes of root diameter and mycorrhizal fungal colonization in 2 temperate grassland communities. New Phytologist 116:129–136

Saize Del Rio JF, Fernandez CE, Bellavita O (1961) Distribution of absorbing capacity of coffee roots determined by radioactive tracers. American Society for Horticultural Science 77:240–244

Schroth G (1998) A review of belowground interactions in agroforestry, focussing on mechanisms and management options. Agroforestry Systems 43:5–34. https://doi.org/10.1007/978-94-017-0679-7_1

Streit J, Meinen C, Rauber R (2019) Intercropping effects on root distribution of eight novel winter faba bean genotypes mixed with winter wheat. Field Crops Research 235:1–10. https://doi.org/10.1016/j.fcr.2019.02.014

Sun B, Gao Y, Yang H, et al (2019) Performance of alfalfa rather than maize stimulates system phosphorus uptake and overyielding of maize/alfalfa intercropping via changes in soil water balance and root morphology and distribution in a light chernozemic soil. Plant and Soil 439:145–161. https://doi.org/10.1007/s11104-018-3888-y

Thorup-Kristensen K, Halberg N, Nicolaisen M, et al (2020) Digging Deeper for Agricultural Resources, the Value of Deep Rooting. Trends in Plant Science 25:406–417. https://doi.org/10.1016/j.tplants.2019.12.007

Thorup-Kristensen K, Rasmussen CR (2015) Identifying new deep-rooted plant species suitable as undersown nitrogen catch crops. Journal of soil and water Conservation 70:399–409. https://doi.org/10.2489/jswc.70.6.399

Wahid PA (2000) A system of classification of woody perennials based on their root activity patterns. Agroforestry Systems 49:123–130

Zhang G, Zhang C, Yang Z, Dong S (2013) Root Distribution and N Acquisition in an Alfalfa and Corn Intercropping System. Journal of Agricultural Science 5:128–142. https://doi.org/10.5539/jas.v5n9p128

